# Nonthermal plasma treatment of polymers modulates biological fouling but can cause material embrittlement

**DOI:** 10.1101/842260

**Authors:** Greg D. Learn, Emerson J. Lai, Emily J. Wilson, Horst A. von Recum

## Abstract

Plasma-based treatment is a prevalent strategy to alter biological response and enhance biomaterial coating quality at the surfaces of biomedical devices and implants, especially polymeric materials. Plasma, an ionized gas, is often thought to have negligible effects on the bulk properties of prosthetic substrates given that it alters the surface chemistry on only the outermost few nanometers of material. However, no studies to date have systematically explored the effects of plasma exposure on both the surface and bulk properties of a biomaterial. This work examines the time-dependent effects of a nonthermal plasma on the surface and bulk properties of polymeric implants, specifically polypropylene surgical meshes and sutures. Findings suggest that plasma exposure improved resistance to fibrinogen adsorption and *Escherichia coli* attachment, and promoted mammalian fibroblast attachment, although increased duration of exposure resulted in a state of diminishing returns. At the same time, it was observed that plasma exposure can be detrimental to the material properties of individual filaments (i.e. sutures), as well as the structural characteristics of knitted meshes, with longer exposures resulting in further embrittlement and larger changes in anisotropic qualities. Though there are few guidelines regarding appropriate mechanical properties of surgical textiles, the results from this investigation imply that there are ultimate exposure limits for plasma-based treatments of polymeric implant materials when structural properties must be preserved, and that the effects of a plasma on a given biomaterial should be examined carefully before translation to a clinical scenario.

## 1. Introduction

Upon implantation, biomedical materials are rapidly covered by proteins, followed by host cells, or bacteria in the case of infection. The nature of these biological attachments often dictate the success of a surgically implanted device. For example, in hernia repair surgery, polymer fabrics called “meshes” are typically used for abdominal wall defect closure. These meshes are commonly associated with complications, particularly post-surgical adhesions and prosthetic infection. Specifically, out of over 350,000 hernia repair procedures performed each year in the United States (Poulose et al., 2012), 13.6% of patients are rehospitalized for adhesion-related complications within 5 years of surgery (Bensley et al., 2013), and between 1-10% of implanted meshes become infected (Falagas and Kasiakou, 2005; Sanchez et al., 2011). Tissue adhesions to mesh are initiated by deposition and maturation of a fibrin matrix on the mesh surface as well as an adjacent tissue/organ (B W J Hellebrekers and Kooistra, 2011), whereas prosthetic infection results from bacterial attachment to the protein-conditioned surface and subsequent biofilm formation. To mitigate such adverse events, many research efforts have focused on altering surface properties of synthetic meshes, either directly or through application of anti-adhesion (Deeken et al., 2012; Schreinemacher et al., 2013) or antibiotic coatings (Badiou et al., 2011; Majumder et al., 2015). However, many synthetic mesh polymers, especially polypropylene (PP), which is the most common mesh material (Coda et al., 2012), are inert, possess low surface energy, and bond poorly (weakly and/or with nonuniform coverage) with coatings.

One strategy that has been explored for direct surface modification, as well as enhancing the uniformity, durability, and adherence of coatings, on difficult substrates like PP is surface treatment with a man-made plasma. Plasma, the fourth state of matter, is composed of a mixture of positive and negative ions, neutral atoms/molecules, radicals, free electrons, and photons (particularly visible and ultraviolet light). It is created when a gas becomes ionized through application of sufficient energy, either through heat (thermal plasma) or electromagnetic fields (nonthermal plasma). Thermal plasmas exist only at high temperatures that would destroy most substrate materials (Desmet et al., 2009), thus only nonthermal plasmas are used for surface modification as considered in this paper.

Interaction of plasma with a polymer surface results in surface cleaning/etching, scission or rearrangement of bonds, and the introduction of new functional groups (as determined by the composition of the carrier gas). In the absence of coatings, these plasma-induced changes can be used to tune biological response to the material. For example, surface engraftment of oxygen can enhance wettability, in turn reducing protein adsorption and conformational denaturation that might otherwise trigger detrimental inflammatory responses on a hydrophobic implant surface (Hu et al., 2001; Lu and Park, 1991), and modulating subsequent cell/bacterial attachment. Alternatively, these plasma-induced changes can contribute to improved receptiveness/adherence to coatings (Navaneetha Pandiyaraj et al., 2015), circumventing the need for hazardous chemicals as coating adhesion promoters, on inert polymers such as PP. Additionally, plasma can easily treat objects with complex geometries and high surface areas. Numerous studies have therefore explored plasma surface treatments for surgical meshes (Avetta et al., 2014; Binnebösel et al., 2012, 2010; Gorgieva et al., 2015; Hu et al., 2017; Junge et al., 2007; Kulaga et al., 2014; Kumar et al., 2013; Labay et al., 2015; Lanzalaco et al., 2019; Lu et al., 2019; Nisticò et al., 2017, 2015, 2013, 2012; Paul et al., 2010; Rivolo et al., 2016; Sanbhal et al., 2019, 2018b; Sannino et al., 2005; Zhang et al., 2014).

Another frequently-argued advantage of plasma-based surface treatment for biomedical polymers such as mesh substrates is that plasma exposure is thought to have negligible effects on bulk mechanical properties, given that the treatment modifies only the outermost few nanometers of the material surface (Chu et al., 2002; Gomathi et al., 2008; Labay et al., 2015; Nisticò et al., 2017, 2015, 2013; Rivolo et al., 2016; Sanbhal et al., 2018b, 2018a; Sannino et al., 2005; Zhang et al., 2014). To our knowledge, however, no prior studies have systematically verified this lack of effect on bulk properties or examined what limits there are to the established beneficial effects of plasma exposure. This paucity of data is concerning for a few reasons: 1) most surgical meshes have relatively high surface-to-volume ratios, 2) plasmas emit photons in the ultraviolet range (Boyd et al., 1997; Jaritz et al., 2017; Liston et al., 1993), 3) ionizing radiation is known to embrittle PP (Abiona and Osinkolu, 2010; Fintzou et al., 2006; Raab et al., 1982; Rabello and White, 1997) and other polymeric materials, and 4) degraded polymer chains at the material surface may resemble nano-scale cracks that could propagate into the bulk of the material upon application of mechanical stress. Considering this, the objective of this work is to evaluate the time-dependent effects of typical nonthermal plasma on both the surface and bulk mechanical properties of surgical mesh devices. We hypothesize that plasma treatment alters biofouling by proteins, cells, and bacteria, but extending exposure accelerates the mechanical failure of PP biomedical textiles.

An overview of this work is depicted in **Figure 1**. PP substrates, primarily surgical textiles and microplates, were treated with nonthermal plasma for selected durations from 0 to 20 minutes. Effects on PP surface composition, protein adsorption, bacterial attachment, and mammalian cell attachment were investigated with X-ray photoelectron spectroscopy (XPS), fluorescence spectroscopy, bioluminescence spectroscopy, and hemacytometry, respectively. Next, effects on mesh material and structural properties were evaluated. Uniaxial tension tests were performed on individual PP filaments and ASTM standard dogbone-shaped mesh specimens cut along two orthogonal axes. Finally, suture retention, tear-resistance, and ball burst tests were performed according to standardized methods (Deeken et al., 2011) using rectangular mesh pieces.

**Figure 1:**
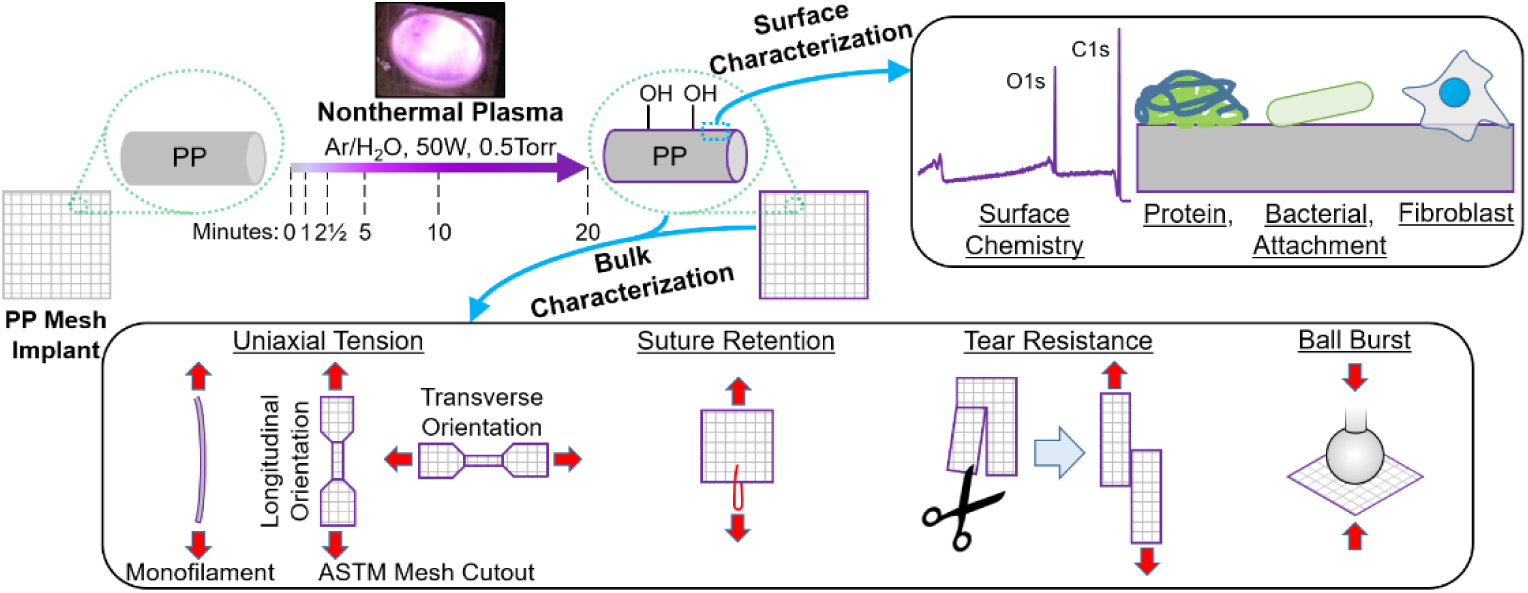
Study overview.

## 2. Materials and Methods

### 2.1 Materials

Prolene PP mesh, Prolene PP Soft mesh, and 4-0 Prolene PP monofilaments were manufactured by Ethicon, Inc. The extruded PP in each of these products is identical in composition according to manufacturer instructions for use. Prolene Soft mesh was used for XPS analysis due to the presence of bundled filaments with wider overall diameters (facilitating higher photoelectron counts), while Prolene mesh and monofilament sutures were used for mechanical tests. The monofilament size of 4-0 was selected to closely approximate the ∼130 µm diameter (Deeken et al., 2011) of filaments in Prolene mesh. For measurement of protein adsorption and bacterial attachment, PP 96-well plates (#M9685) were purchased from Sigma Aldrich, fluorescein isothiocyanate (FITC)-labeled fibrinogen (#FIB-FITC) was purchased from Molecular Innovations, Luria Bertani (LB) broth (#BP1426-500) and tissue culture polystyrene 96-well plates (#07-200-90) were purchased from Fisher Scientific, and bioluminescent ilux pGEX(-) *Escherichia coli* was purchased from Addgene, a gift from Dr. Stefan Hell (plasmid #107879) (Gregor et al., 2018). For measurement of fibroblast attachment, PP 24-well plates (#1185U58) and lids (#1185U62) were purchased from Thomas Scientific, tissue culture polystyrene 24-well plates (#08-772-1H), DMEM-LG (#10-567-014), fetal bovine serum (#16-141-079), penicillin-streptomycin (#15-140-122) were purchased from Fisher Scientific, and NIH/3T3 fibroblasts (#AKR-214) were purchased from Cell BioLabs, Inc.

### 2.2 Plasma Treatment

PP materials were exposed to plasma at varying durations to examine the effects of treatment time on both surface and the bulk properties. For studies on surface chemistry and mechanical properties, meshes and monofilaments were removed from sterile packaging, cut to size, and pre-treated for 1, 2.5, 5, 10, or 20 minutes with a low-pressure 500 mTorr (67 Pa) Ar/H_2_O nonthermal plasma (50 W, 13.56 MHz) using a Branson/IPC Model #1005-248 Gas Plasma Cleaner. Plasma treatment was performed within 18 h of XPS analysis and 6 h of mechanical testing. For protein, bacterial, and mammalian cell attachment experiments, PP 96-well and PP 24-well plates were plasma-treated in similar fashion within 2 h of biofoulant exposure. Non-treated (0 min) PP samples with no known prior exposure to ultraviolet light were included as controls in all experiments, and tissue culture polystyrene (a common substrate for cell culture and attachment *in vitro*) microplate surfaces were included in protein, bacterial, and mammalian cell attachment experiments.

### 2.3 Surface Characterization

Effects of plasma exposure on PP surfaces were evaluated in terms of surface composition, protein adsorption, bacterial attachment, and mammalian cell attachment. These were investigated using XPS, fluorescence spectroscopy, bioluminescence spectroscopy, and hemacytometry, respectively.

#### 2.3.1 X-ray Photoelectron Spectroscopy

Effects of plasma exposure on PP implant surface chemistry were determined using XPS. Prolene Soft mesh surfaces were analyzed for elemental content using a PHI Versaprobe 5000 Scanning X-Ray Photoelectron Spectrometer equipped with Al Kα source (h? = 1486.6 eV). Scans were acquired on 1 scan location per sample per plasma treatment duration. Survey scans were collected using a 200 µm spot size, 45 W power, 15 kV acceleration voltage, 117.40 eV pass energy, 0.40 eV step size, 25 ms/step, 8 cycles, 44.7° take-off angle, and 0-1100 eV range. The C1s peak was auto-shifted to 284.8 eV, and the ratios of the elements carbon, nitrogen, oxygen, and fluorine were analyzed. The areas of peaks were taken with background set using a Shirley function from 280-292 eV for C1s, 396-404 eV for N1s, 526-538 eV for O1s, and 675-695 eV for F1s. After survey scans, high-resolution scans were collected using a 100 µm spot size, 25.2 W power, 15 kV acceleration voltage, 23.50 eV pass energy, 0.20 eV step size, 50 ms/step, 16 cycles, 44.7° take-off angle, and 278-298 eV range for C1s. The vertical sampling depth ζ for take-off angle θ = 44.7° and reported inelastic mean free path of λ = 3.5nm at a photoelectron kinetic energy of 1 keV for PP surfaces (Cumpson, 2001), is estimated to be ∼7.4 nm based on the relation (Ton-That et al., 2000) ζ = 3λcos(θ). Analysis was performed using MultiPak software version 9.8.0.19 (Physical Electronics, Inc.).

#### 2.3.2 Fibrinogen Adsorption

Protein adsorption is one of the first stages in biological fouling occurring after implantation. While many proteins adsorb and compete for an implant surface, fibrinogen is one of the most studied, and was chosen as a model protein here given its common involvement in inflammatory responses to biomaterials (Hu et al., 2001; Zdolsek et al., 2007) and post-surgical adhesion formation (B. W J Hellebrekers and Kooistra, 2011). FITC-labeled fibrinogen was thawed and diluted to a working concentration of 50 µg/mL in sterile PBS containing 2 mM sodium azide. This solution was pipetted onto PP 96-well plates at 100 μL/well before covering plates with Parafilm and foil, and incubating them overnight at 37°C. Following incubation, all wells were rinsed with sterile PBS such that aspiration was performed 5 times, standard curves with known FITC-fibrinogen concentrations were pipetted in empty rows for each microplate, and a Biotek Synergy H1 plate reader was used to read the fluorescence in each well using excitation and emission wavelengths of 490 nm and 525 nm, respectively. Data presented reflects 12 wells/condition from one experiment, and the same trends were confirmed across one additional independent experiment (not shown).

#### 2.3.3 Bacterial Attachment

Attachment of bacteria to a biomaterial surface is a key step that precedes any prosthetic infection, and *E. coli* was chosen as a model microorganism for bacterial attachment to PP given its common involvement in mesh infection following repair of incarcerated abdominal hernias (Yang et al., 2015), being derived from the gut. Bioluminescent *E. coli* were thawed from frozen stock, inoculated into a 14 mL round-bottom tube of sterile LB broth with vented lid, and expanded in suspension for 18 h in a dedicated 37°C incubator with the tube lid vented. The tube was then removed from the incubator and stored at 4°C to maintain bacteria in the stationary phase. Prior to seeding onto experimental surfaces, the bacterial suspension was diluted to an optical density at 600 nm (OD600) of 0.50 relative to sterile LB broth. The bacterial suspension was then directly seeded onto PP 96-well plates at 100 µL/well, cultured under static conditions for 24 h at 37°C, and following incubation, wells were rinsed 4 times with 200 µL/well sterile PBS, then emptied and filled a final time with 100 µL/well sterile LB broth. Several known dilutions of bacteria were included for creation of standard curves in duplicate for each type of surface. Bioluminescence measurement was then performed using a Biotek Synergy H1 plate reader (luminescence endpoint scan, 5 s integration, 1 mm read height, full light emission, 135 gain, top optics, 100 ms delay, extended dynamic range). Data presented reflects 14-16 wells/condition from one experiment, and the same trends were observed across one additional independent experiment using PP mesh substrates (not shown).

#### 2.3.4 Fibroblast Attachment

Attachment of host cells to a biomaterial surface is an important event that can lead to tissue ingrowth around the implant as adherent cells deposit and remodel matrix proteins. Fibroblasts were chosen as a model mammalian cell type given their role in incorporation of mesh implants with host abdominal wall tissue, a step that necessary to anchor the mesh in place and prevent its migration. NIH/3T3 fibroblasts were cultivated on 100 mm tissue culture polystyrene dishes in culture medium consisting of 89% DMEM high glucose, 10% fetal bovine serum, and 1% penicillin-streptomycin, and cells were subcultured before reaching confluence. Fibroblasts were trypsinized, resuspended, counted, and seeded at 30,000 cells/cm^2^ onto PP 24-well plates at 500 μL/well, then allowed to attach for 18h in a 37°C incubator under 5% CO_2_ and 95% humidity. To wash away non-adherent cells, each well was gently rinsed thrice with 750 μL sterile PBS. Next, 200 μL/well trypsin, followed by two rinses with 350 μL/well culture medium, were used to detach cells and collected for cell quantification using a hemacytometer. Data presented reflects 6 wells/condition from one experiment, with one hemacytometry count performed for each well.

### 2.4 Bulk Characterization

A variety of mechanical tests were performed to determine the extent of bulk property changes following plasma treatments. These tests included uniaxial tension for individual PP filaments and dogbone-shaped mesh specimens, and suture retention, tear-resistance, and ball burst tests of rectangular mesh specimens. Unless otherwise specified, tests were performed using an Instru-Met renewed load frame (#1130) operated using Testworks 4 software, an Instron 100 lbf (445 N) tension load cell (cat #2512-103), and a sampling rate of 100 Hz. Failure load was defined as the maximum load (N) sustained by the specimen during the test, which for some tests may not have coincided with failure displacement. Failure displacement was defined as the magnitude of crosshead travel (mm) at which the specimen completely ruptured and could no longer bear load. Work to failure was defined as the energy (J) absorbed by the specimen from the beginning of the test (0 mm) up to the failure displacement.

#### 2.4.1 Uniaxial Tension Testing of PP Monofilaments

PP monofilament sutures were tested in uniaxial tension to examine effects of plasma exposure on mechanical properties of mesh implants at the level of their most basic component – the individual filaments. A total of three Prolene 4-0 monofilaments were removed from sterile packaging and cut in half, with each half being assigned to a 0 min or 20 min plasma treatment (n = 3 samples/condition). Each monofilament half was then cut into 7 pieces (technical replicates) of 2.5” (63.5 mm) length, such that a total of 21 pieces were tested per treatment group. Pieces were gripped on each end to a depth of 0.75” (19 mm) to leave a gauge length of 1” (25.4 mm), and tested in uniaxial tension at 10 mm/min until failure. An Instron 10 lbf (44.5 N) tension load cell (cat #2512-111) and a sampling rate of 100 Hz were used. Data presented reflects 3 samples/condition.

#### 2.4.2 Uniaxial Tension Testing of PP Meshes

Unlike individual filaments, meshes tend to exhibit a complex geometric structure, so were tested in uniaxial tension to examine effects of plasma exposure on mechanical properties along two orthogonal axes. Methods for uniaxial tension testing were adapted from Deeken *et al*. (Deeken et al., 2011), with slight modification. Meshes were die-cut into ASTM D412 Type A dogbone shaped samples along both the longitudinal and transverse directions. Samples were gripped on each end to a depth of 1” (25.4 mm) to leave a gauge length of 3.5” (88.9 mm). Uniaxial tension testing was performed at 50 mm/min until mesh failure. Data presented reflects 4-7 longitudinal mesh samples/condition, and 4-5 transverse samples/condition.

#### 2.4.3 Suture Retention Testing of PP Meshes

Suture retention testing was performed to simulate loading of plasma-treated meshes through a single fixation point or “suture” that passes through it, which in this case is represented by a stainless steel wire to ensure that failure is localized only within the mesh structure. Methods for suture retention testing were adapted from Deeken *et al*. (Deeken et al., 2011). A custom test fixture was machined that consisted of a 0.014” (0.36 mm) diameter stainless steel wire that could be clamped in place on an aluminum frame, after passing the wire through a mesh 1 cm from its bottom edge. Mesh specimens were cut to 1” × 2” (25.4 mm × 50.8 mm). Meshes were installed with a gauge length of 1” (25.4 mm) and clamped at the top end using pneumatic grips. Each mesh specimen was tested in tension at a rate of 300 mm/min until the suture pulled through the mesh. The “pull-out force” was defined as the failure load (N). A sampling rate of 200 Hz was used. Meshes were oriented such that the direction of pull was parallel to the longitudinal axis. Data presented reflects 4-10 mesh samples/condition.

#### 2.4.4 Tear Resistance Testing of PP Meshes

Tear resistance testing was performed to simulate the ability of a plasma-treated mesh to resist further tearing once a tear has already begun. Methods for tear resistance testing were adapted from Deeken *et al*. (Deeken et al., 2011). Specimens were cut to 1” × 2” (25.4 mm × 50.8 mm). A 1” (25.4 mm) slit was cut parallel to the longitudinal axis from the middle of the short edge toward the center of the mesh to form 2 tabs or “pant legs.” The left and right tabs were clamped to a ½” (12.7 mm) depth in the upper and lower grips, respectively. This arrangement yielded a 1” (25.4 mm) gauge length. The test was conducted at a rate of 300 mm/min until the specimen tore in half. The “tear force” was defined as the failure load (N). A sampling rate of 200 Hz was used. Meshes were oriented such that the direction of pull was parallel to the mesh longitudinal axis. Data presented reflects 6-8 mesh samples/condition.

#### 2.4.5 Ball Burst Testing of PP Meshes

Ball burst testing was performed to simulate a loading scenario which plasma-treated meshes could be anticipated to experience in clinical use (e.g. during coughing or sneezing), specifically that of pressure applied perpendicular to the mesh plane. Methods for ball burst testing were adapted from Deeken *et al*. (Deeken et al., 2011) and ASTM D3787-16 (ASTM International, 2016). Fixtures were machined that consisted of a polished stainless steel ball having a 1” (25.4 mm) diameter for the top fixture, and an aluminum ring clamp having a 1.75” (44.5 mm) internal diameter with neoprene-lined faces to prevent mesh slippage for the bottom fixture. Specimens were cut to 2.75” × 2.75” (70 mm × 70 mm). The steel ball was pressed transversely through the mesh at a rate of 300 mm/min until the mesh burst. The “burst force” was defined as the failure load (N). Circumference at failure refers to the circumference (cm) of the contact zone that the mesh made with the steel ball at the instant of failure and was determined using Solidworks 2017 Student Edition (Dassault Systèmes SolidWorks Corporation), as calculated using the displacement value at which the burst force was reached. “Burst strength” refers to the maximum spherical wall tension sustained by the mesh and was calculated by dividing the burst force by the circumference at failure. A Material Testing System MTS 810 frame (MTS Systems Corporation) was used along with a 2000 lbf (8900 N) load cell. Data presented reflects 5-6 mesh samples/condition.

### 2.5 Statistical Analysis

All data is presented as mean ± standard deviation. Statistical analysis tests were carried out in Microsoft Excel 2016 using two-sample two-tailed Student’s t-tests with unequal variance, with statistical significance being set at p < α = 0.05. A Bonferroni correction was applied in fibrinogen adsorption and bacterial attachment experiments to set statistical significance at p-values less than α divided by 15, the number of Student’s t-test comparisons performed among the 6 conditions. Data points in any individual group data set that fell more than 2 standard deviations away from the mean of that data set were considered outliers and excluded from analyses.

## 3. Results

### 3.1 Surface Characterization

Plasma-induced effects on PP material surfaces were evaluated in terms of surface chemistry, and attachment of proteins, bacteria, and mammalian cells.

#### 3.1.1 X-ray Photoelectron Spectroscopy of PP Meshes

XPS was used to follow the extent of changes in PP surface chemistry in response to plasma exposure time. Analysis of XPS survey scan spectra (**Fig 2**) indicated that plasma enhanced the atomic percentage of oxygen on PP substrate surfaces in a duration-dependent manner (**Fig 3**) with little apparent impact on surface nitrogen content. For the untreated substrate, trace amounts of fluorine were detected on the mesh surface, possibly derived from device packaging or the textile production process. The high-resolution C1s spectrum was nearly symmetric for the untreated mesh surface, while spectra were skewed left following plasma treatment for any duration (**Fig 4**). High-resolution spectra also showed an increasing signal at ∼289 eV with longer plasma exposures, attributed to carboxylic acids and/or carbonates on the surface, species that can only be produced through PP chain scission.

**Figure 2:**
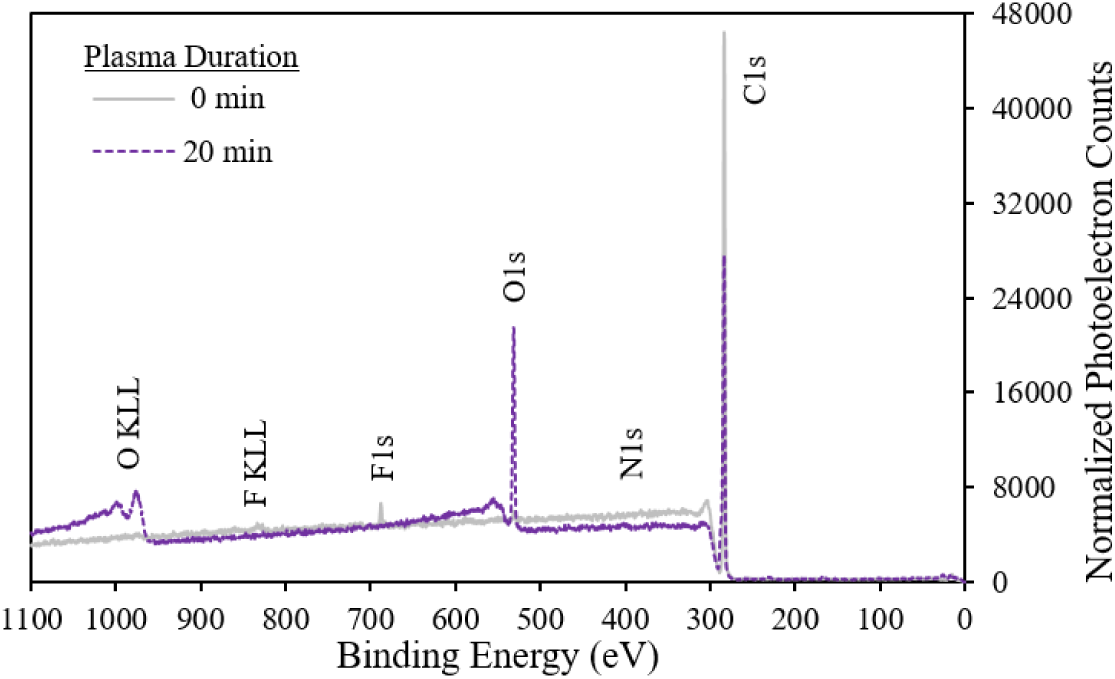
XPS survey scans, normalized by area under the curve, for 0 min and 20 min plasma treatment durations on Prolene Soft mesh. KLL peaks indicate atomic relaxation via the Auger effect, and were not used for analysis.

**Figure 3:**
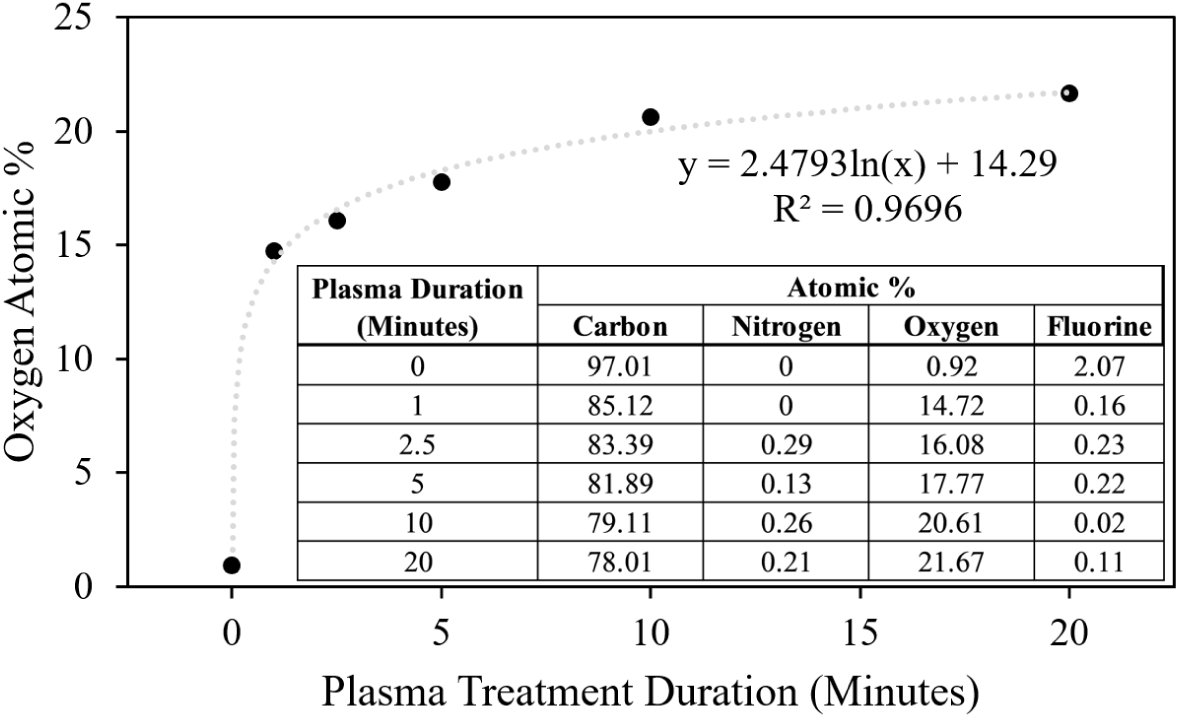
Quantification of XPS survey scans to determine the atomic composition of mesh surfaces (n = 1 mesh/group). Data is fit with an exponential growth curve.

**Figure 4:**
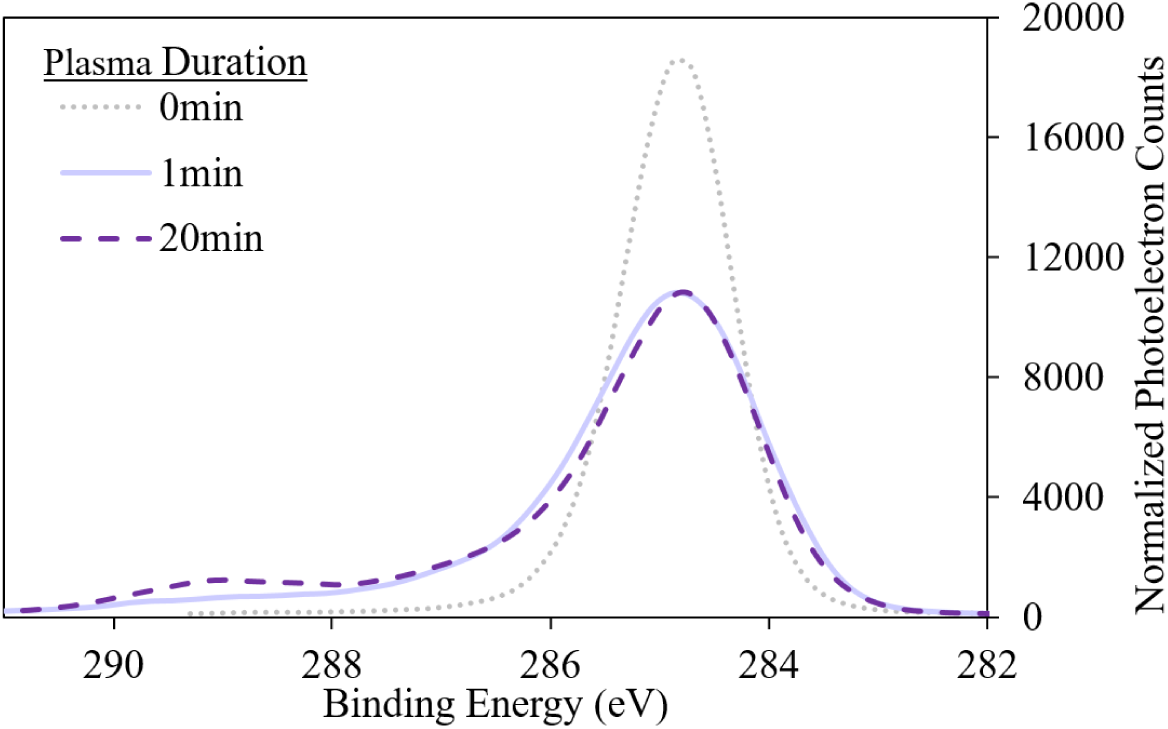
XPS high-res C1s scans, normalized by area under the curve, demonstrated increasing counts at ∼289 eV with increasing treatment duration.

#### 3.1.2 Fibrinogen Adsorption

Fibrinogen was used as a model protein to evaluate changes in protein adsorption to PP surfaces following plasma exposure. Plasma treatment for any length of time reduced fibrinogen adsorption by roughly half (p < 0.001) relative to untreated PP (**Fig 5**). Increasing the length of plasma treatment time beyond 1 min resulted in diminishing resistance to fibrinogen adsorption (p < 0.002). For reference, the average rate of fibrinogen adsorption to tissue culture polystyrene in the same experiment was 10.2 ± 0.8%.

**Figure 5:**
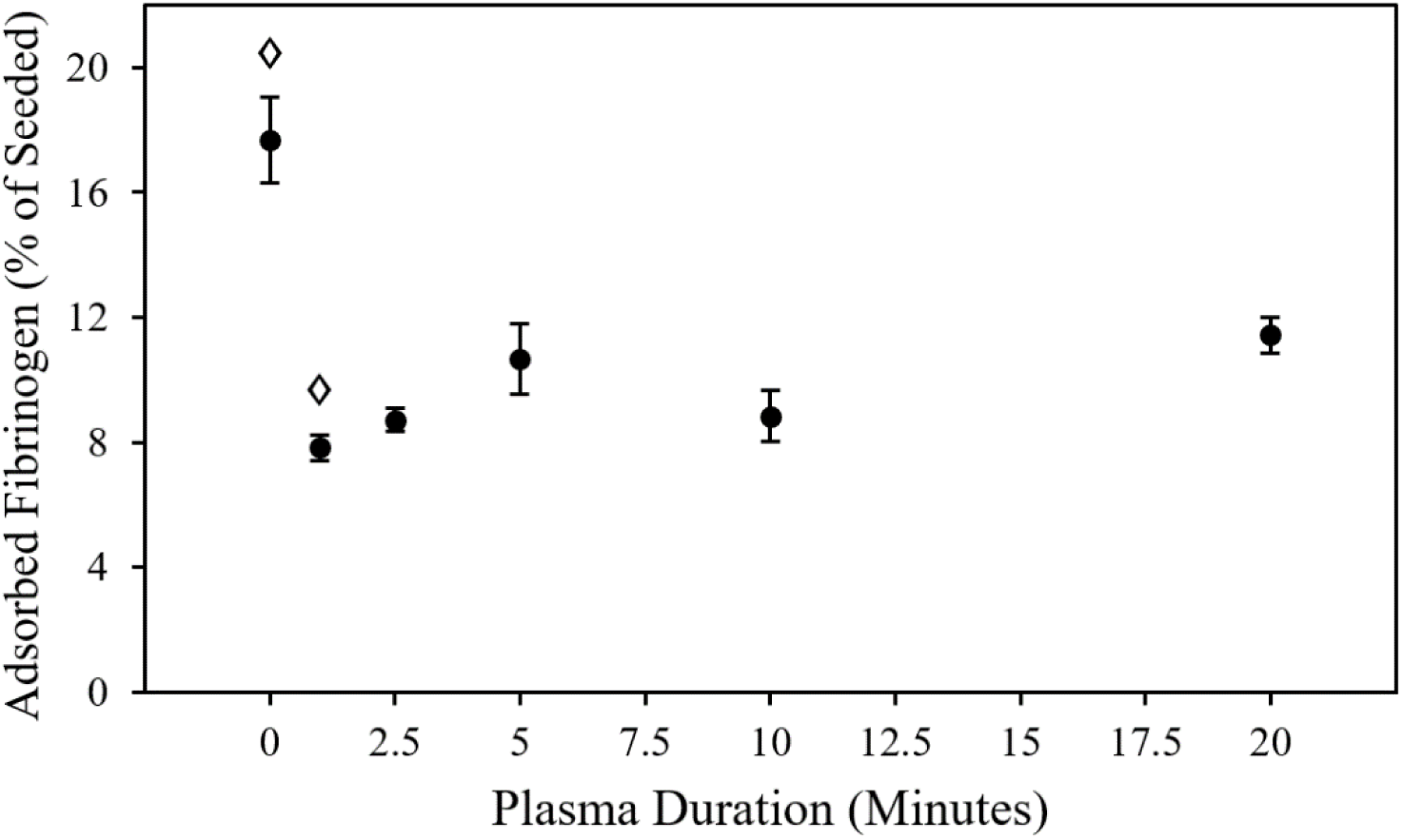
Effect of plasma treatment duration on FITC-fibrinogen protein adsorption. ◊Significant difference to all other time points.

#### 3.1.3 Bacterial Attachment

Materials were exposed to *E. coli* to evaluate the capacity for plasma treatment to reduce bacterial attachment to PP surfaces. Plasma treatment for any length of time reduced attachment of *E. coli* (p < 0.001) by a proportion of at least 75% relative to untreated PP (**Fig 6**). Bacterial attachment appeared to level off after 1 min of plasma treatment (p > 0.048), until attachment rates of roughly half that seen at 1 min (p < 0.001) were seen at both 10 min and 20 min of plasma treatment. For reference, the average rate of *E. coli* attachment to tissue culture polystyrene in the same experiment was 2.36 ± 0.59%.

**Figure 6:**
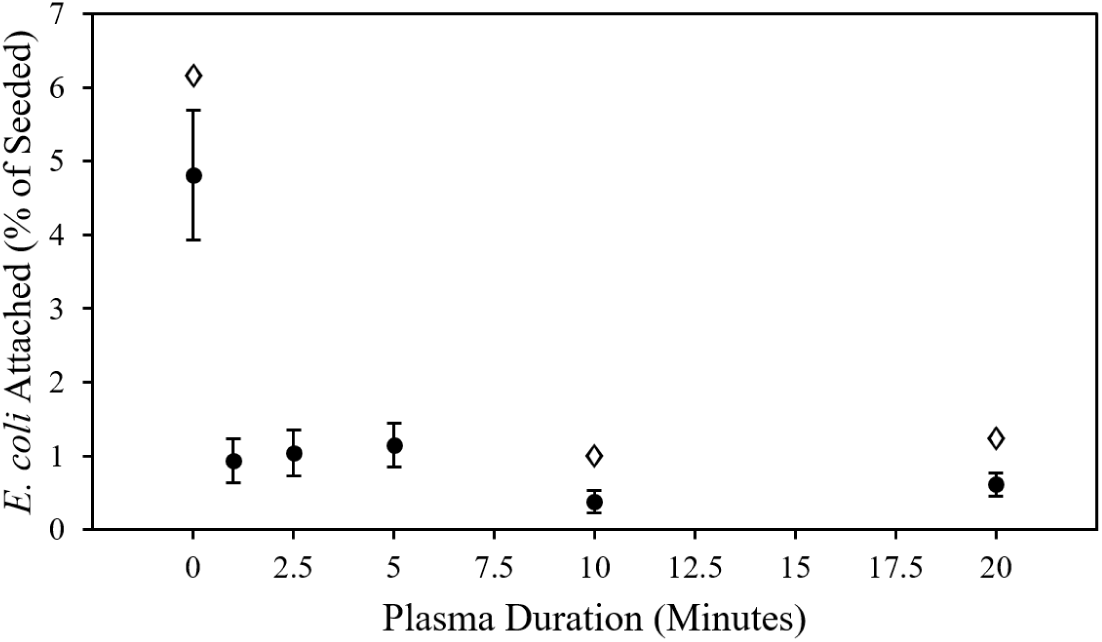
Effect of plasma treatment duration on *E. coli* bacterial attachment. ◊Significant difference to all other time points.

#### 3.1.4 Mammalian Fibroblast Attachment

Substrates were exposed to fibroblasts to evaluate the capacity for plasma treatment to enhance mammalian cell attachment to PP surfaces. Plasma treatment for any length of time increased attachment of NIH/3T3 fibroblasts (p < 0.018) relative to untreated PP (**Fig 7**). Fibroblast attachment to untreated PP was not detectable. Increasing the length of plasma treatment time beyond 1 min did not result in any significant improvements in fibroblast attachment (p > 0.1). For reference, the average rate of NIH/3T3 attachment to tissue culture polystyrene in the same experiment was 64.3 ± 7.4%.

**Figure 7:**
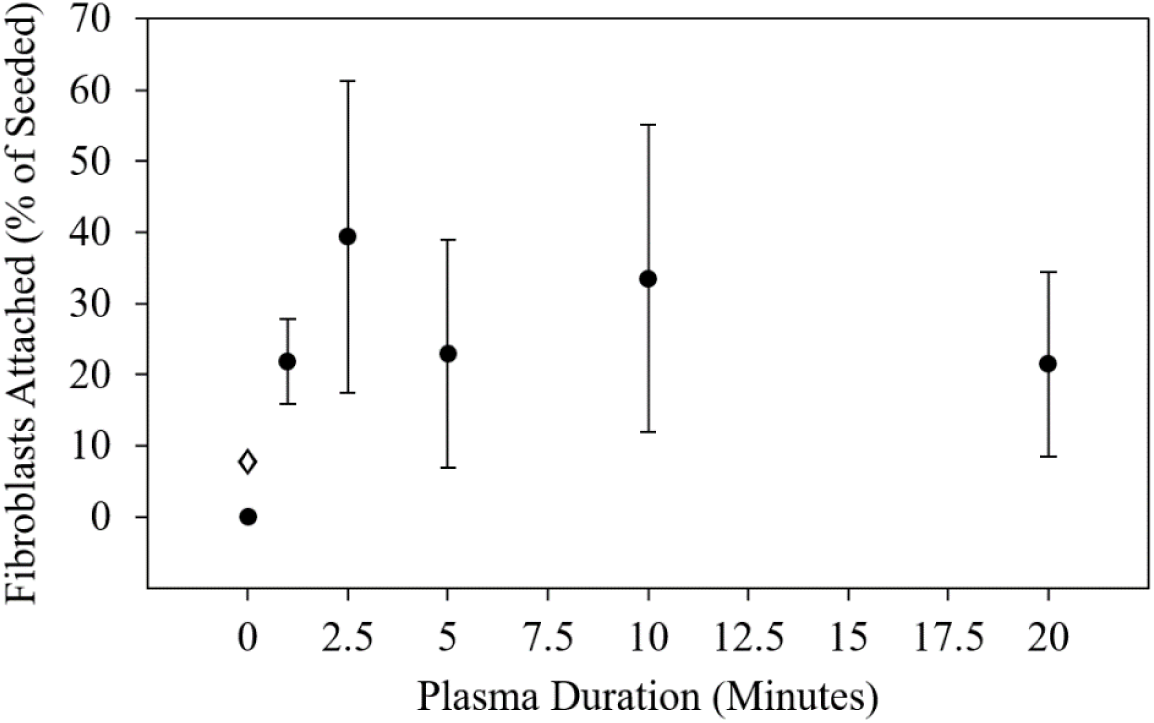
Effect of plasma treatment duration on mammalian fibroblast attachment. ◊Significant difference to all other time points.

### 3.2 Bulk Characterization

A variety of mechanical tests were performed to assess plasma-induced bulk property changes of PP biomedical textiles. The tests used for evaluating these effects on textile implant structural properties are depicted in **Fig 8**.

**Figure 8:**
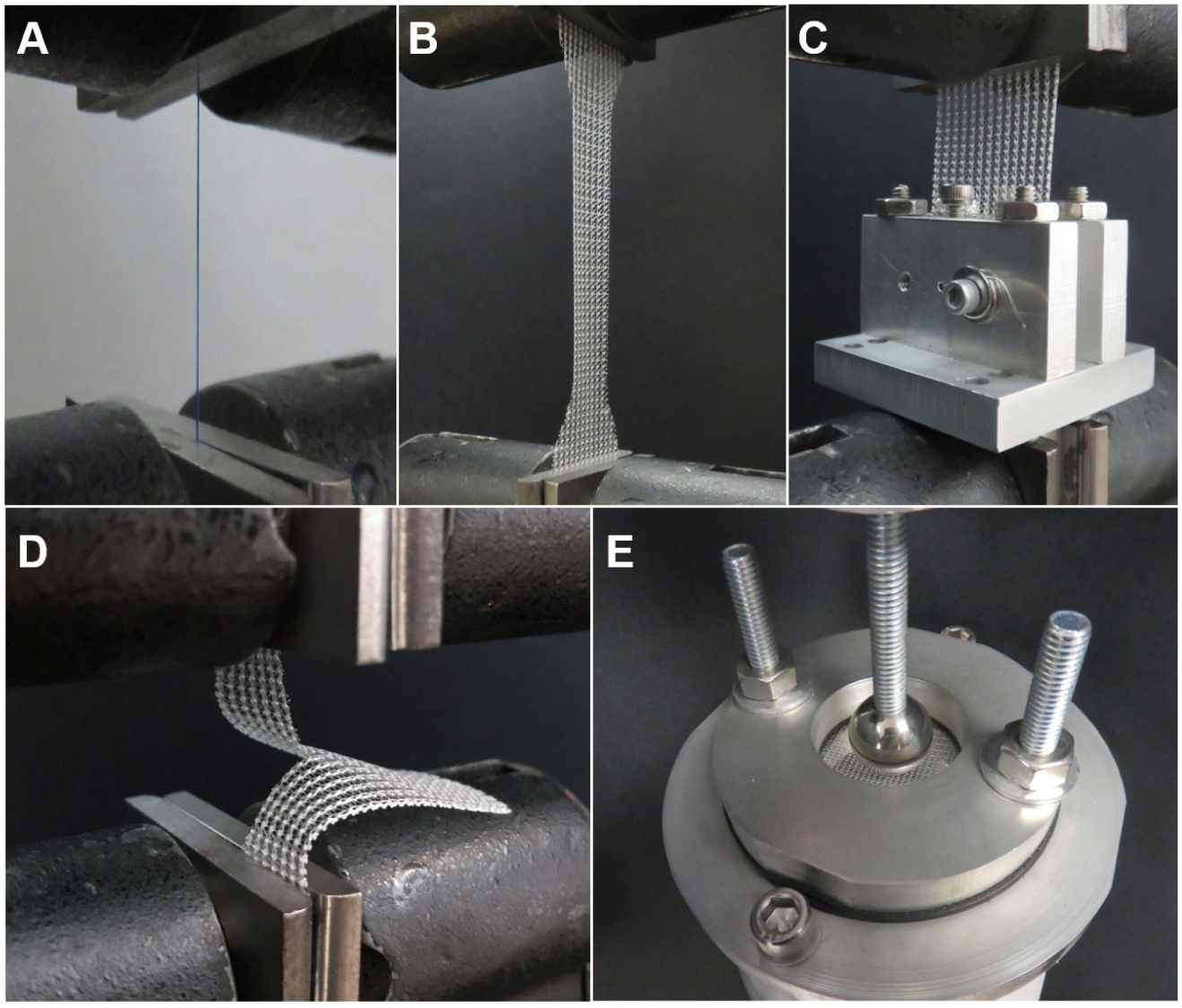
Tests performed to assess the effects of plasma on material and structural properties of PP surgical meshes: (a) uniaxial tension tests on PP monofilaments, and (b) uniaxial tension, (c) suture retention, (d) tear-resistance, and (e) ball burst tests on PP meshes.

#### 3.2.1 PP Monofilament Uniaxial Tension Testing

PP monofilament sutures were tested in tension to explore the effects of plasma exposure on bulk property changes of mesh implants at the level of their simplest component. Plasma treatment for 20 min caused an 11% decrease in maximum load (p = 0.046), a 43% decline in work to failure (p = 0.01), and a 32% reduction in displacement to failure (p = 0.013) for Prolene monofilaments (**Table 1** and **Fig 9**).

**Table 1:** Effects of plasma exposure time on PP monofilament tensile properties. *Significant difference to untreated.

| Plasma Duration (min) | Monofilament Uniaxial Tension |  |  |
| --- | --- | --- | --- |
|  | Load to Fail (N) | Work to Fail (mJ) | Displacement to Fail (mm) |
| 0 | $16.70 \pm 0.22$ | $234.79 \pm 18.62$ | $20.09 \pm 1.41$ |
| 20 | $14.80 \pm 0.80^*$ | $133.13 \pm 28.42^*$ | $13.72 \pm 1.96^*$ |

**Figure 9:**
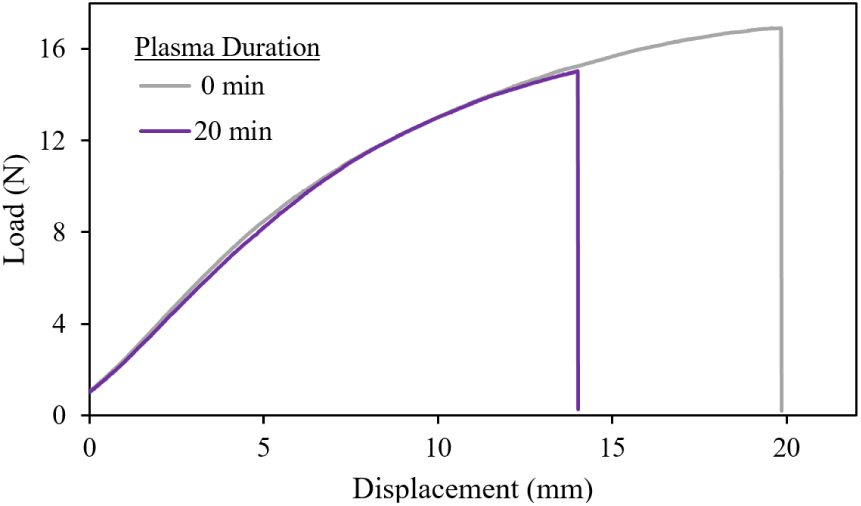
Plasma treatment resulted in embrittlement of Prolene 4-0 monofilaments, as depicted by representative (closest to average values) load-displacement curves.

#### 3.2.2 PP Mesh Uniaxial Tension Testing

Meshes were tested in uniaxial tension to evaluate effects of plasma exposure on textile implant mechanical properties along two orthogonal axes (**Fig 10**), referred to as longitudinal and transverse. Plasma treatment decreased failure load (p < 0.001), work to failure (p < 0.006), and displacement to failure (p < 0.001) of Prolene hernia mesh material in the longitudinal direction (**Fig 11a** and **Table 2**). Longer treatments were associated with weaker, more brittle meshes. For example, plasma treatment for 20 min caused a 41% decrease in max load, a 27% decrease in failure displacement, and a 54% decrease in work to failure. While the declines in failure displacement are comparable, these declines for max load and work to failure are proportionally greater than the decreases observed for monofilaments treated for the same length of time. Additionally, with the exception of a ∼5% decrease in displacement to failure at all time points (p < 0.03) and a ∼12% decrease in work to failure at 20 min (p = 0.048), there were otherwise no apparent effects of plasma on Prolene mesh max load (p > 0.3) and work to failure (p > 0.1) in the transverse direction (**Fig 11b** and **Table 2**). Altogether, these results suggest that plasma-induced effects on textile bulk mechanics are influenced by structure (e.g. the geometry and orientation of the knit) in addition to material properties, with effects being more drastic for the predominant load-bearing axis.

**Table 2:**
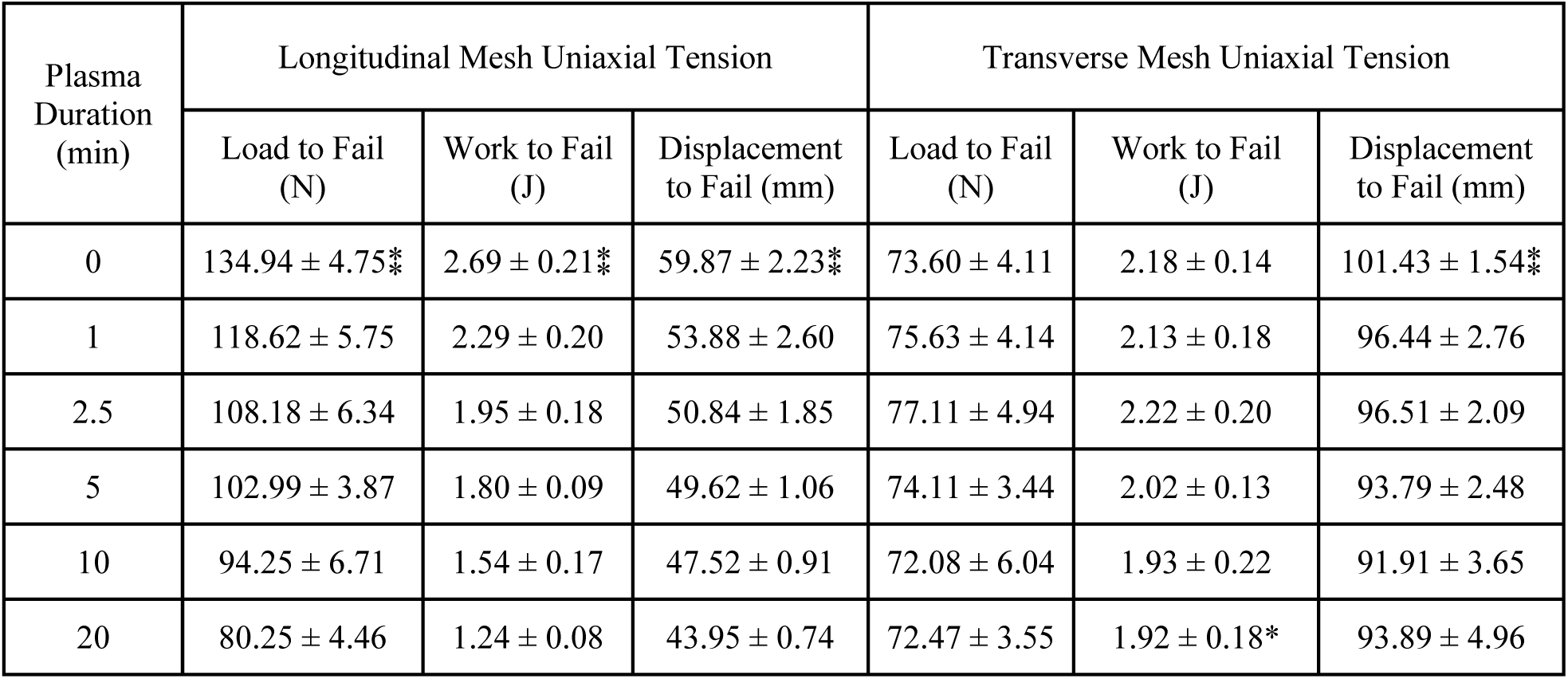
Effects of plasma exposure time on mesh longitudinal and transverse mechanics. *Significant difference to untreated. ⁑Significant difference to all subsequent time points.

| Plasma Duration (min) | Longitudinal Mesh Uniaxial Tension |  |  | Transverse Mesh Uniaxial Tension |  |  |
| --- | --- | --- | --- | --- | --- | --- |
|  | Load to Fail (N) | Work to Fail (J) | Displacement to Fail (mm) | Load to Fail (N) | Work to Fail (J) | Displacement to Fail (mm) |
| 0 | 134.94 ± 4.75* | 2.69 ± 0.21* | 59.87 ± 2.23* | 73.60 ± 4.11 | 2.18 ± 0.14 | 101.43 ± 1.54* |
| 1 | 118.62 ± 5.75 | 2.29 ± 0.20 | 53.88 ± 2.60 | 75.63 ± 4.14 | 2.13 ± 0.18 | 96.44 ± 2.76 |
| 2.5 | 108.18 ± 6.34 | 1.95 ± 0.18 | 50.84 ± 1.85 | 77.11 ± 4.94 | 2.22 ± 0.20 | 96.51 ± 2.09 |
| 5 | 102.99 ± 3.87 | 1.80 ± 0.09 | 49.62 ± 1.06 | 74.11 ± 3.44 | 2.02 ± 0.13 | 93.79 ± 2.48 |
| 10 | 94.25 ± 6.71 | 1.54 ± 0.17 | 47.52 ± 0.91 | 72.08 ± 6.04 | 1.93 ± 0.22 | 91.91 ± 3.65 |
| 20 | 80.25 ± 4.46 | 1.24 ± 0.08 | 43.95 ± 0.74 | 72.47 ± 3.55 | 1.92 ± 0.18* | 93.89 ± 4.96 |

**Figure 10:**
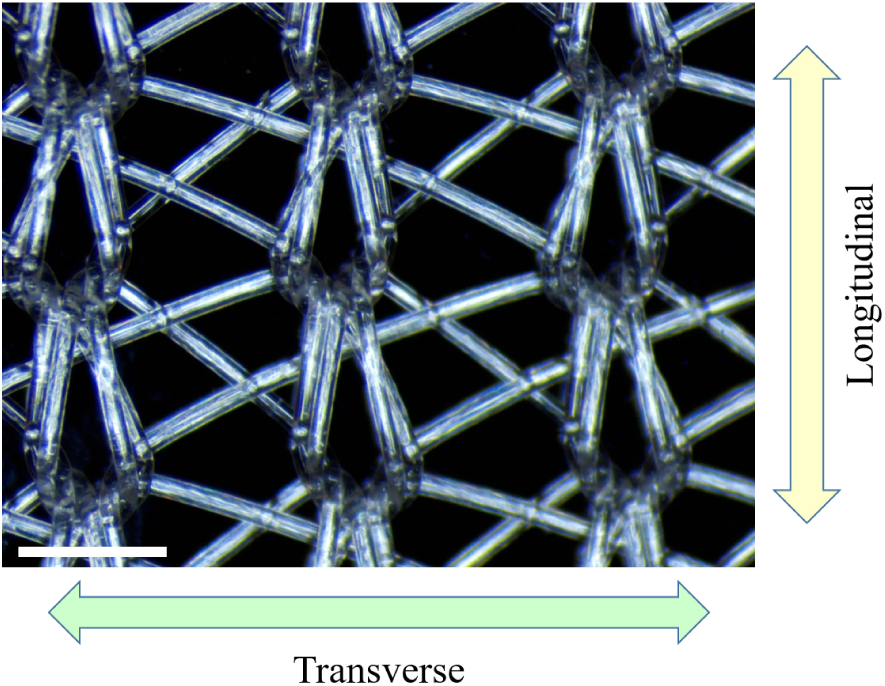
Longitudinal and transverse orientations, defined as being parallel to the direction of crosshead travel for uniaxial tension tests on Prolene meshes. Scale bar = 1 mm.

**Figure 11:**
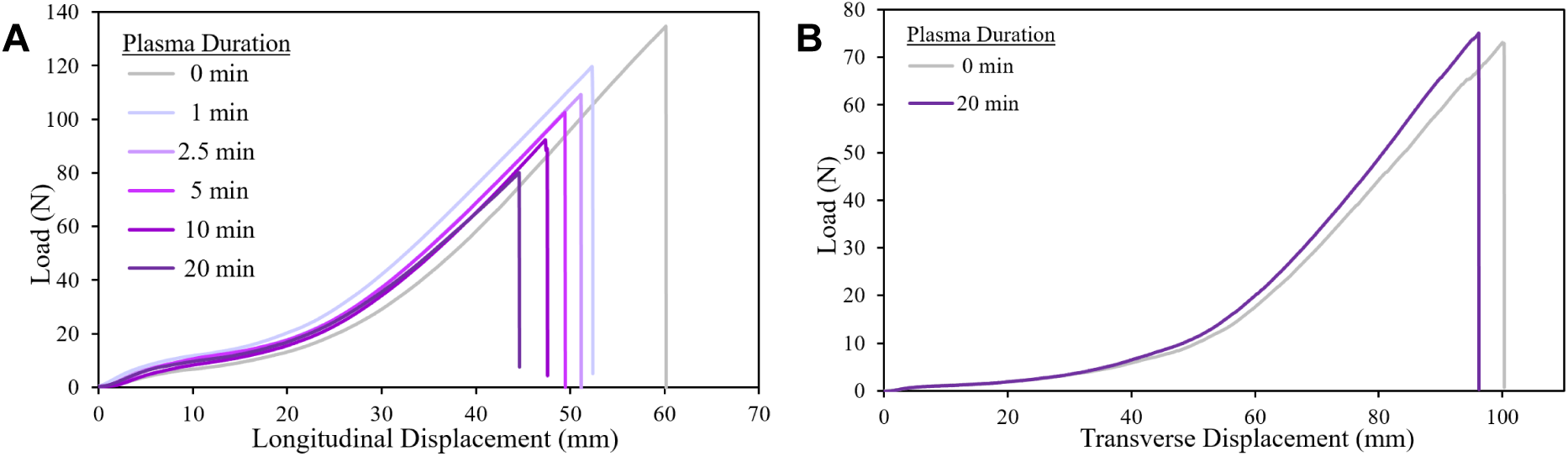
Representative load-displacement curves from uniaxial tension tests of Prolene mesh in the longitudinal (a) and transverse (b) directions.

#### 3.2.3 Mesh Suture Retention Testing

Suture retention testing was conducted to examine the impact of plasma treatment on the ability of mesh samples to withstand loading through a single fixation point. Plasma treatment for 20 min was observed to result in a 17% reduction in pull-out force (p = 0.006) compared to untreated meshes (**Fig 12**). No other treatment duration showed a significant difference to untreated meshes in pull-out force (p > 0.08). Note that displacement and work to failure are not considered precise metrics in this test, given that the wire may have captured different numbers of filaments to break through on different samples, depending on how the mesh was cut and where in the knit pattern the wire was inserted. The longitudinal suture retention strength of 58.9±7.5 N measured in this study for untreated Prolene is in agreement with the 61.2 N previously reported in literature (Deeken and Lake, 2017).

**Figure 12:**
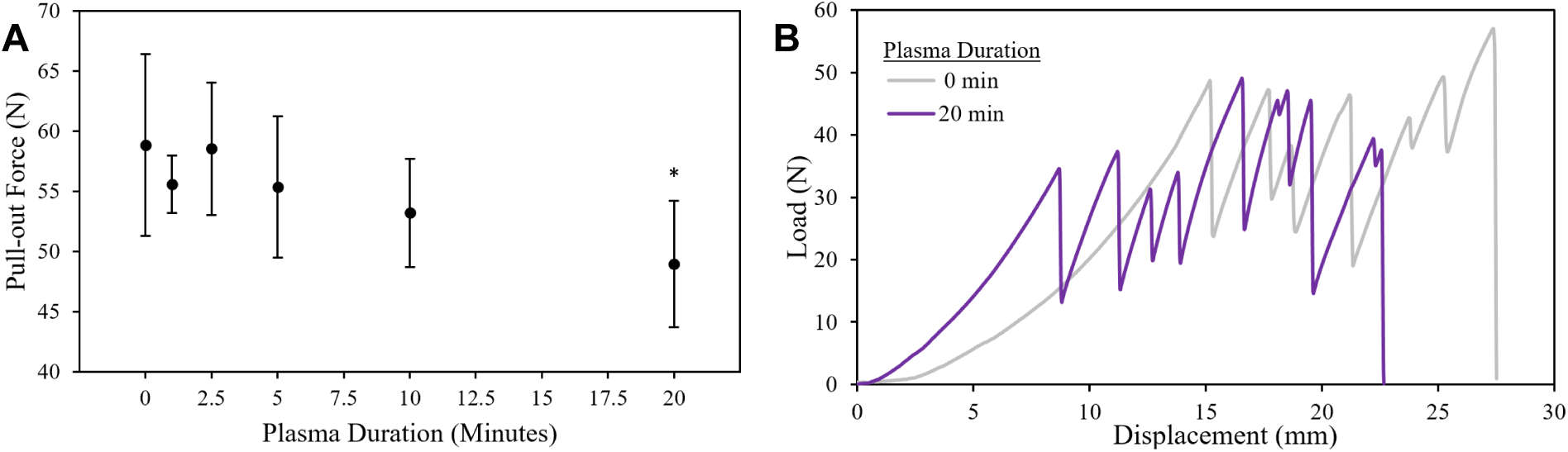
Mesh suture retention tests in the longitudinal direction showing effect of plasma duration on maximum pull-out force (a), and representative load-displacement curves (b). *Represents significant difference to untreated.

#### 3.2.4 Mesh Tear-Resistance Testing

Tear-resistance testing was carried out to explore the effect of plasma treatment on the resistance of mesh samples to tear propagation once a tear has already been initiated. Plasma treatment had no discernible impact (p > 0.2) on mesh tear force (**Fig 13**). Representative load-displacement curves are shown in **Fig 13b**. Results from tear-resistance testing demonstrated high variability in terms of tear force due to the lack of consistency in how the tear propagated through the mesh among different trials. For example, it was observed that the tear may propagate and exit through either the side (long edge) or center (short edge) of the rectangular specimen. These tear propagation directions are equivalent to those seen in the longitudinal and transverse tensile tests, respectively, given that during tensile tests the tear propagates perpendicular to the direction of pull. Interestingly, a higher proportion of side failures were observed for plasma-treated meshes at all time points than untreated meshes (**Table 3**), and this is consistent with the finding that plasma treatments preferentially embrittled meshes tested in the longitudinal direction. At any rate, attempting to account for the different tear propagation patterns would reduce sample size by different proportions among groups, therefore any effects of plasma treatment on tear force may be obscured by this variability. The longitudinal tear force of 27.8±12.8 N measured in this study for untreated Prolene is in reasonable agreement with the 33.66 N previously reported in literature (Deeken and Lake, 2017).

**Table 3:** Failure patterns for tear-resistance testing among samples after plasma treatments.

| Plasma Duration (min) | n | Side Failures | Center Failures |
| --- | --- | --- | --- |
| 0 | 7 | 1 | 6 |
| 1 | 7 | 5 | 2 |
| 2.5 | 7 | 4 | 3 |
| 5 | 6 | 4 | 2 |
| 10 | 6 | 4 | 2 |
| 20 | 8 | 5 | 3 |

**Figure 13:**
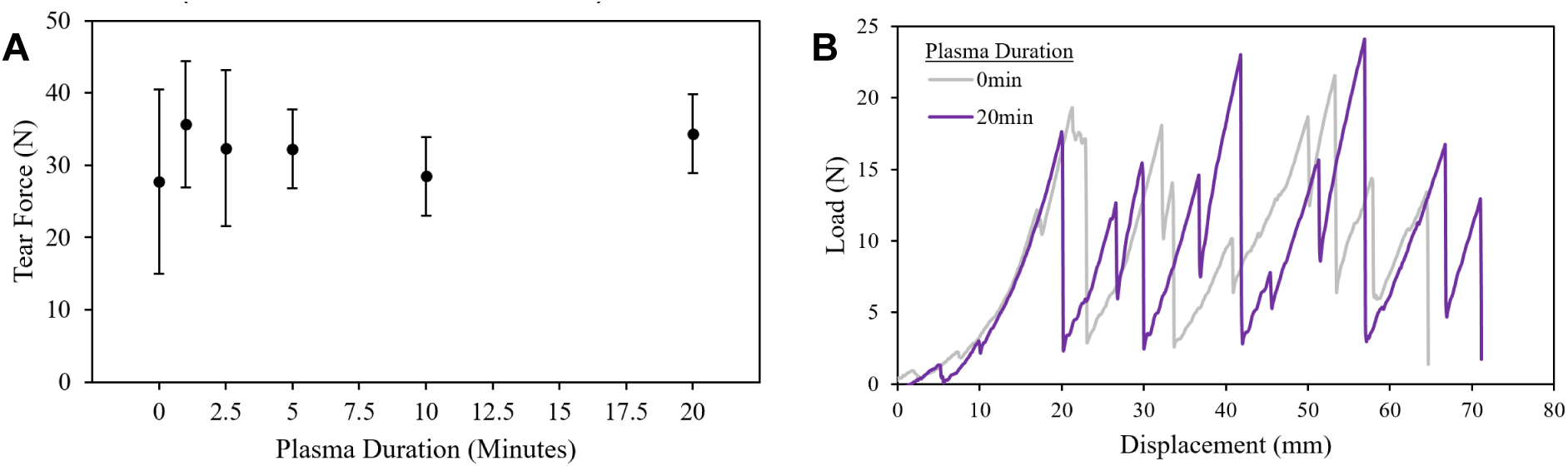
Effects of plasma treatment duration on maximum load sustained in PP mesh tear-resistance tests (a). Representative load-displacement curves (b).

#### 3.2.5 Mesh Ball Burst Testing

Ball burst testing was conducted to examine the impact of plasma treatment on the ability of meshes to withstand biaxial loading that might be encountered during changes in intra-abdominal pressure *in vivo*. For ball burst tests, plasma treatments for 10 and 20 min caused significant declines in burst force (p < 0.006), displacement to failure (p < 0.02), and work to failure (p < 0.02) compared to untreated (**Fig 14** and **Table 4**). In particular, after 20 min of treatment, burst force diminished by 19%, work to failure decreased by 39%, and displacement to failure declined by 13%. Interestingly, this proportional decrease in work to failure is roughly equal to that seen for Prolene monofilaments. No significant differences to untreated meshes were observed for earlier time points (p > 0.09). The burst strength measured in this study for untreated Prolene is slightly less than the 156.6 N previously reported in literature (Deeken and Lake, 2017). This discrepancy could perhaps be due to minor procedural differences in determination of displacement and subsequent calculation of circumference at burst.

**Table 4:** Effects of plasma treatment duration on ball burst properties. *Significant difference to untreated.

| Plasma Duration (min) | Burst Force (N) | Work to Fail (J) | Displacement to Fail (mm) | Circumference at Burst (cm) | Burst Strength (N/cm) |
| --- | --- | --- | --- | --- | --- |
| 0 | 847.03 ± 72.62 | 6.31 ± 1.30 | 23.06 ± 1.57 | 6.62 ± 0.21 | 127.71 ± 7.76 |
| 1 | 838.78 ± 100.38 | 6.05 ± 1.12 | 22.93 ± 1.16 | 6.61 ± 0.16 | 126.66 ± 12.47 |
| 2.5 | 796.21 ± 82.78 | 5.28 ± 1.04 | 21.89 ± 1.14 | 6.45 ± 0.19 | 123.21 ± 9.33 |
| 5 | 773.19 ± 86.70 | 5.04 ± 0.93 | 21.56 ± 1.12 | 6.42 ± 0.17 | 120.22 ± 10.24 |
| 10 | 718.89 ± 28.43* | 4.44 ± 0.33* | 20.80 ± 0.47* | 6.30 ± 0.07* | 114.03 ± 3.50* |
| 20 | 689.21 ± 30.32* | 3.86 ± 0.28* | 20.05 ± 0.42* | 6.18 ± 0.07* | 111.40 ± 3.58* |

**Figure 14:**
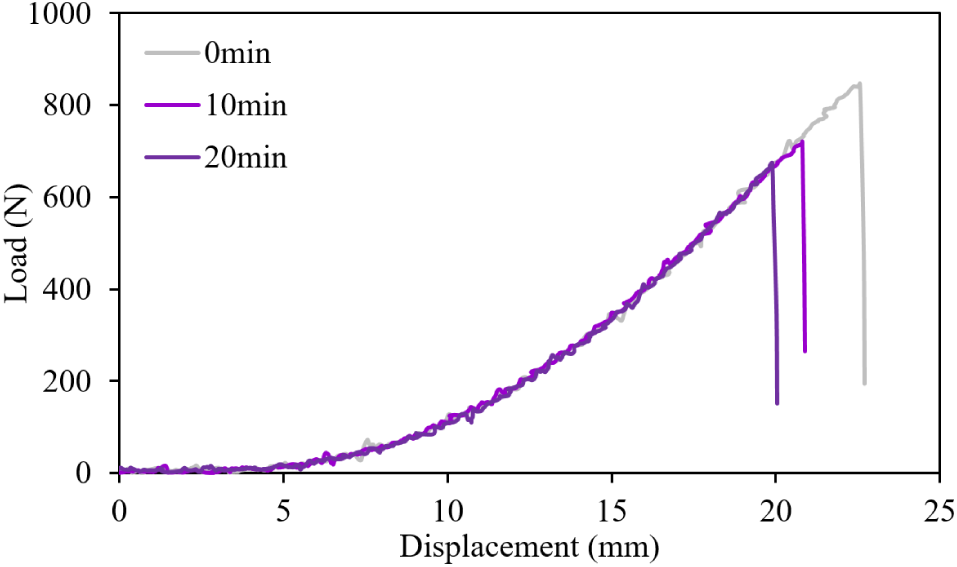
Representative load-displacement curves for ball-burst tests.

## 4. Discussion

These experiments were designed to test the hypothesis that plasma treatment alters biological fouling by proteins, cells, and bacteria, but increasing the duration of exposure accelerates the mechanical failure of PP biomedical textiles. Findings regarding fibrinogen, fibroblast, and *E. coli* attachment, as well as findings from uniaxial tension tests of monofilaments (sutures), and longitudinal tension, suture retention, and ball burst tests of meshes support this hypothesis. Effects of plasma duration on mesh transverse tension and tear resistance properties were less severe, however both of these tests emphasized failure in the axis of the mesh that bears less load. We can state with certainty, however, that while there were beneficial effects on protein, cell, and bacterial attachment, the nonthermal plasma in this study had non-negligible effects on the bulk properties of biomedical textiles. Additionally, the plasma treatment parameters used here are similar to those of plasmas applied towards surface modification of biomaterials currently used clinically, or being investigated for such use. Power outputs between 40 W and 200 W, treatment durations between 30 s and 10 min, and carrier gases such as Ar, N_2_, O_2_, and NH_3_ have been reported for plasma treatments of surgical meshes (Hu et al., 2017; Lanzalaco et al., 2019; Nisticò et al., 2015; Rivolo et al., 2016; Sanbhal et al., 2019, 2018b; Sannino et al., 2005; Zhang et al., 2014).

There are several possible explanations for the demonstrated embrittlement effect of plasma on PP. As demonstrated by XPS high-resolution C1s scans, longer plasma treatments appeared to result in an increase in the proportion of carboxylic acids and carbonates on the PP surface. Considering that the tertiary carbon atom in the PP repeat unit has the highest radical stability (hence is the most likely reaction site), these oxygen-rich functional groups can only be created through some degree of chain scission at the polymer surface. These degraded molecular chains at the surface may act like nano-scale cracks, increasing in size with longer plasma exposure, that could propagate into the bulk of the material upon application of sufficient mechanical stress. In support of this reasoning, prior studies have demonstrated that plasma treatments roughen the topology of polymer surfaces (Sanbhal et al., 2018).

There have been several studies on hernia mesh mechanical properties (Corey R Deeken et al., 2011; Corey R. Deeken et al., 2011; Eliason et al., 2011; Est et al., 2017; Pott et al., 2012). Though there are few definitive guidelines for the mechanical properties that a surgical mesh should possess, it is important to be aware of the detrimental effects that plasma treatments may have on hernia mesh bulk mechanical properties, and to try to minimize these through careful selection of plasma processing parameters. Based on our results, tension tests along the predominant load-bearing direction and ball burst tests would appear to be the most sensitive assays at detecting detrimental effects of plasma treatment on a given mesh fabric. The guidelines that exist regarding mesh failure properties recommend a suture retention strength that exceeds 20 N and a maximum burst pressure over 50 N/cm (Deeken and Lake, 2017). Although plasma treatments for up to 20 min did not result in embrittlement of Prolene mesh below any of these particular failure values, a PP mesh having lower baseline mechanical properties (e.g. Prolene Soft mesh) or higher surface-to-volume ratio could potentially have been weakened to such an extent.

A notable finding from this study was that plasma treatments of PP for all durations tested significantly reduced fibrinogen adsorption and *E. coli* attachment relative to untreated surfaces, and increasing exposure time had diminishing returns on this passive repellence. Likewise, exposure to plasma for any length of time significantly enhanced fibroblast attachment relative to untreated surfaces, while increasing plasma duration had no additive effect on this attachment. Hydrophobic materials such as untreated PP have a strong tendency to adsorb and unfold (i.e. denature) protein solutes. It is energetically favorable for proteins to displace water molecules repelled by a hydrophobic surface and then change conformation such that core nonpolar segments can associate tightly with the surface via nonpolar interactions. Proteins adsorbed on the material then serve as a conditioning layer requisite for further biofouling events, such as bacterial or cell attachment. Plasma treatment renders the surface more hydrophilic, thereby altering the concentrations and conformations of the adsorbed proteins, and modulating subsequent biofouling events. It has previously been shown that plasma treatment decreases protein adsorption of both albumin and fibrinogen to PP (Navaneetha Pandiyaraj et al., 2015). But to our knowledge, no studies have assessed the effects of plasma exposure time on protein, bacterial, or cell resistance of PP. Our findings regarding protein and bacterial resistance of plasma-treated PP suggest that plasma treatment may be a strategy to reduce the likelihood or severity of adhesion formation and prosthetic infection on PP mesh surfaces. At the same time, our findings regarding fibroblast attachment on plasma-treated PP suggest that plasma treatment may promote mesh incorporation with the abdominal wall. Finally, our findings suggest that long exposures (with their associated decline in bulk mechanical properties) are not necessary to realize these beneficial effects.

This study sets the stage for several possible additional investigations. Future studies are needed to determine whether the effects of plasma seen in these studies on PP meshes extend to meshes that exhibit different geometries, or meshes made from other polymeric materials such as polyester or poly(tetrafluoroethylene). In particular, meshes that exhibit a higher surface-to-volume ratio, or meshes that are more lightweight and flexible at baseline should be considered, as plasma-induced embrittlement of these could pose more dangerous consequences. Prolene mesh is a relatively rigid, heavyweight mesh with high baseline mechanical properties (Deeken et al., 2011), and it is considered to maintain its strength indefinitely in clinical use according to manufacturer product literature. It was chosen as a representative mesh for several reasons: 1) its homogenous composition of monofilament PP, the most common mesh material (Coda et al., 2012), 2) its longstanding clinical use in abdominal surgery for over 4 decades, and 3) its constituent Prolene material is used in several Ethicon surgical products (including Prolene sutures, Prolene Soft mesh, and Proceed composite mesh) that span a wide variety of surgical purposes. Additional studies on plasma-treated meshes should also explore fatigue properties, which are more physiologically relevant than monotonic tests given the periodic nature of loading *in vivo* (e.g. breathing, coughing, jumping, etc.), to evaluate whether plasma treatments predispose meshes to failure after repetitive movements. Further investigations should also consider the effects of plasma-treated mesh storage time on bulk properties, as hydrophobic recovery of plasma-treated PP surfaces normally occurs over several weeks in the absence of a coating to stabilize the activated surface (Jokinen et al., 2012). Finally, the effects of plasma treatment on tissue adhesion formation and prosthetic infection of surgical textiles could also be investigated *in vivo*.

## 5. Conclusions

These experiments sought to systematically quantify the positive and negative effects of nonthermal plasma on polymeric implants. While plasma treatment for any length of time increased surface oxygen content, and reduced fibrinogen adsorption and *E. coli* attachment on PP surfaces, increasing the length of exposure resulted in diminishing benefit with regards to surface chemistry, protein and bacterial resistance, and mammalian cell attachment. Simultaneously, plasma treatments were found to result in bulk embrittlement of PP biomedical textiles across several standardized testing modalities, with the duration of exposure being correlated to the decline in mechanical properties. Taken together, the results from this study indicate that plasma-based treatments of PP surgical meshes should be optimized for both surface qualities and implant structural properties (if they are of chief importance), and the effects of a plasma process on a given biomedical material at both the surface and bulk levels should be carefully evaluated before translation to any clinical scenario.

## Acknowledgments

The authors acknowledge support through National Institutes of Health: NIH R01 GM121477 (HvR), and NIH Ruth L. Kirschstein NRSA T32 AR007505 Training Program in Musculoskeletal Research (GDL). Additional support was provided by the Center for Stem Cell and Regenerative Medicine Undergraduate Student Summer Program (ENGAGE) at Case Western Reserve University (EJL). Valuable core facility services were provided by the Swagelok Center for Surface Analysis of Materials, the Advanced Manufacturing and Mechanical Reliability Center, and Think[box] at CWRU. The authors also thank Kevin Abbassi for expertise with XPS, Chris Tuma for expert assistance with mechanical testing, Katherine Yan for technical help, and Nathan Rohner and Alan Dogan for revision suggestions.

